# In silico generation and augmentation of regulatory variants from massively parallel reporter assay using conditional variational autoencoder

**DOI:** 10.1101/2024.06.25.600715

**Authors:** Weijia Jin, Yi Xia, Sai Ritesh Thela, Yunlong Liu, Li Chen

## Abstract

Predicting the functional consequences of genetic variants in non-coding regions is a challenging problem. Massively parallel reporter assays (MPRAs), which are an *in vitro* high-throughput method, can simultaneously test thousands of variants by evaluating the existence of allele specific regulatory activity. Nevertheless, the identified labelled variants by MPRAs, which shows differential allelic regulatory effects on the gene expression are usually limited to the scale of hundreds, limiting their potential to be used as the training set for achieving a robust genome-wide prediction. To address the limitation, we propose a deep generative model, MpraVAE, to *in silico* generate and augment the training sample size of labelled variants. By benchmarking on several MPRA datasets, we demonstrate that MpraVAE significantly improves the prediction performance for MPRA regulatory variants compared to the baseline method, conventional data augmentation approaches as well as existing variant scoring methods. Taking autoimmune diseases as one example, we apply MpraVAE to perform a genome-wide prediction of regulatory variants and find that predicted regulatory variants are more enriched than background variants in enhancers, active histone marks, open chromatin regions in immune-related cell types, and chromatin states associated with promoter, enhancer activity and binding sites of cMyC and Pol II that regulate gene expression. Importantly, predicted regulatory variants are found to link immune-related genes by leveraging chromatin loop and accessible chromatin, demonstrating the importance of MpraVAE in genetic and gene discovery for complex traits.

## Introduction

Noncoding variants play a crucial role in gene regulation by being located in cis-regulatory elements such as enhancer and promoter, where they affect the binding of transcriptional factor, dysregulate mRNA splicing and impact the 3D chromatin conformation [1, 2]. Due to these significant biological functions, noncoding variants have been implicated in various diseases. For example, TERT promoter mutations are known to play a fundamental and widespread role in tumorigenesis [3]. Additionally, a risk variant rs6983267 in transcriptional enhancer demonstrates long-range interaction with MYC in colorectal cancer [4]. Pathogenic splice-site mutations can cause complete skipping of CFTR exon 7, leading to a severe cystic fibrosis phenotype. Despite their important role, discovering causal noncoding variants remains a challenging task in human genetics research. Although genome wide association studies (GWAS) and expression quantitative trait loci (eQTL) analyses have identified tens of thousands of putative causal variants in noncoding regions associated with human diseases and traits, linkage disequilibrium (LD) complicates pinpointing the exact causal variants.

Alternatively, massively parallel reporter assays (MPRAs), which are an *in vitro* high-throughput method, can simultaneously test thousands of variants by evaluating the existence of allele specific regulatory activity. This capacility allows MPRAs to perform high-throughput functional screen of SNPs and small insertions/deletions within the tight LD region of GWAS or eQTLs loci, facilitating the identification of causal variants. Consequently, MPRA experiments can generate high-quality labelled variants that indicate the regulatory potential across diseases, traits and cell lines. Nevertheless, despite the capacity of MPRAs to scale up the number of tested variants to tens of thousands, the identified labelled variants, which shows differential allelic regulatory effects on the gene expression, are usually limited to hundreds. This scale is still small compared to an approximately 600 million variants identified in gnomAD [5]. Moreover, it is experimentally infeasible to evaluate all 600 million variants due to technical difficulty and financial cost. Therefore, only a small set of GWAS and eQTL loci are carefully selected for MPRA experiments. The full landscape of functional variants in the entire human genome remains elusive.

With the development of sequencing technologies, multi-dimensional functional annotations have become available through public consortiums such as ENCODE and Roadmap Epigenomics Project [6, 7]. These enriched data resources have enabled the development of computational methods for predicting genome-wide functional variants. These computational methods can be classified into unsupervised or supervised methods. Unsupervised methods, such as FunSeq2, adopt a weighted scoring method to integrate conservation score, transcription factor binding events, and enhancer-gene linkages directly [8] . Other unsupervised methods, such as fitCons, FitCons2 and DVAR, leverage the evolutionary evidence to assess the fitness consequences of variants [9–11]. Supervised methods, on the other hand, use functional annotations such as the tissue and cell type-specific open chromatin, transcription factor binding and histone modification derived from ENCODE and Roadmap Epigenomics as features for labelled variants. These labeled variants are collected from resources such as Human Gene Mutation Database (HGMD), ClinVar, NHGRI GWAS Catalog (https://www.ebi.ac.uk/gwas/) and COSMIC [12–14]. For example, FATHMM-XF and FATHMM-MKL utilizes HGMD, ncER adopts both ClinVar and HGMD, and CScape uses COSMIC [15–18]. These supervised methods also vary in terms of prediction models used: FATHMM-XF, FATHMM-MKL, and CADD adopt SVM, while ncER utlizes XGBoost.

Though these methods have been widely used for predicting noncoding functional variants, their prediction are often general because the variants in ClinVar, HGMD and NHGRI GWAS Catalog comes from heterogenous diseases, traits and tissues. In contrast, MPRA experiments target specific diseases or traits and are profiled by specific cell lines, which generate labelled variants with a disease- or trait-specific regulatory effect. Some approaches, such as DIVAN, TIVAN and WEVar, have been developed to predict disease-specific risk variants and tissue-specific regulatory variants [19–23]. However, these approaches rely on statistically significant disease-specific GWAS SNPs and tissue-specific cis-eQTLs as the surrogates for the causal variants in their training sets, which may lead to biased prediction.

Ideally, MPRA variants, directly validated by wet-lab experiments, should be used as the training set to develop a supervised machine learning model for context-specific prediction. Nevertheless, the limitations of throughput of MPRA experiment, including low throughput, high cost, and labor-intensive protocol, result in few labelled variants being available. This scarcity poses a challenge for adopting classic supervised machine learning methods, which typically require large training sample sizes to achieve robust prediction.

Data augmentation has been widely adopted in image analysis to improve the performance and robustness of the prediction models by artificially expanding the training data through generating synthetic data [24–26]. Adding synthetic datasets to the training process helps overcome the scarcity of the training data, increase data variability, prevent potential overfitting and enhance feature learning. Simple data augmentation operations leverage basic image manipulations such as cropping, rotation, flipping and scaling. Recent development in deep learning technique, especially deep generative models, have led to more sophisticated data augmentation methods. Among these, generative adversarial network (GAN) and variational autoencoder (VAE) are two of the most popular approaches. GAN consists of a generator and a discriminator: the generator produces synthetic data from random noise, while the discriminator distinguishes between both real and synthetic data. The adversarial training process pits the two components against each other, and onced trained, the generator can produce synthetic data that closely resembles the real data. Similarly, VAE consists of an encoder and a decoder. The encode maps the data into a latent space, typically characterized by a probabilistic distribution defined by mean and variance. The decoder reconstructs the data from the samples generated from the distribution, minimizing the difference between reconstruction loss and Kullback-Leibler (KL) divergence to learn the model parameters. Once trained, the encoder can generate synthetic data resembling real data from the learnt distribution. The advantage of GAN and VAE over simple data augmentation methods is their ability to learn augmentation directly from the data, enabling more complex and meaningful transformations. These models can generate highly realistic augmentations that better reflect real-world variations and scenarios. For instance, they have improved the phenotype prediction from microbiome data and enhanced cell type identification and feature learning from single cell data [27–30].

Motivated by the widely acclaimed success of deep generative models used in data augmentation, we propose a novel deep generative model-based data augmentation method, named MpraVAE, paired with a convolutional neural network, to improve the functional prediction of noncoding variants. MpraVAE offers several key contributions: (i) enhanced training data: As a data augmentation method, MpraVAE can generate synthetic labeled variants that closely resemble labeled MPRA variants, thereby augmenting the training set and improving prediction performance. To the best of our knowledge, MpraVAE is the first deep generative model specifically developed for this purpose. (ii) innovative variant representation: MpraVAE converts the task of variant augmentation into DNA sequence augmentation by representing the one-hot encoded flanking sequences of variants as a 2D image. This approach allows the use of deep generative models for data augmentation. (iii) class-specific generation: MpraVAE incorporates the categorical labels of MPRA variants, indicating whether or not they have allelic regulatory effects, by using a conditional variational autoencoder to generate class-specific variant representations; (iv) advanced loss function: MpraVAE adopts a novel reconstruction loss function that maximizes the similarity between real and synthetic sequences at both the single-base level and the k-mer level; (v) user-friendly web server: an RShiny web server hosts the pretrained MpraVAE models, allowing users to upload variants and obtain prediction scores easily. By benchmarking MpraVAE on several MPRA datasets, we demonstrate that it achieves substantially better prediction performance compared to baseline method, existing data augmentation methods and outperform numerous functional annotation methods. We further extend the application of MpraVAE to identify genome-wide regulatory variants and explore their distribution patterns. Moreover, we explore the SNP-gene link by leveraging chromatin loop and accessible chromatin to validate the findings in a case study.

## Results

### Overview of MpraVAE

The overall goal of our data augmentation method is to *in silico* generate synthetic MPRA variants, which will boost the sample size in the model training and, in turn, improves the prediction performance. Instead of creating synthetic MPRA variants directly, our proposed MpraVAE, a deep generative model, generates synthetic flanking DNA sequences that closely resemble the real flanking DNA sequences of observed MPRA variants. MpraVAE has two key considerations. First, since MPRA variants are labelled into positive and negative sets, data generation needs to consider the class label of variants. Second, the loss function, which defines the similarity between observed and synthetic flanking sequence, needs to be carefully designed to account for biological sequence properties. To address these considerations, we develop MpraVAE, employing a Conditional Variational Autoencoder (CVAE) architecture to generate synthetic sequences that resemble the flanking DNA sequences from both positive and negative MPRA variants derived from real MPRA experiments (**Figure 1**). MpraVAE consists of a conditional encoder *φ* and a conditional decoder *θ*. The encoder *φ* takes both the one-hot encoding sequence *X*, representing the variant, and the label for the variant *Y* (1 for positive and 0 for negative) as input to learn the conditional latent distribution *q*(*Z*|*X*,*Y*) of the latent variable *Z* given the sequence *X* and label *Y*. *q*(*Z*|*X*,*Y*) is essentially a normal distribution *N*(*μ*(*X*,*Y*),*σ*^2^(*X,Y*)) with parameters are estimated from both *X* and *Y.* The decoder takes both *Z* and *Y* to generate the reconstructed one-hot encoding sequence denoted as *X̂*. To learn the network parameters for both encoder *φ* and decoder *θ*, MpraVAE minimizes the total loss, which is a weighted sum of three individual losses. The first and the second losses are reconstruction loss, designed to ensure the synthetic sequences resemble the real sequences. The first loss is the mean difference of the one-hot encoding sequence, which encourages similarity between synthetic and real sequences on the base level. The second loss is k-mer loss, which is the mean difference of k-mer frequency, ensuring similarity at the k-mer level by accounting for spatially correlated nucleotides that play important functional roles. The third loss is the KL-divergence loss, which ensures the learnt latent distribution is close to the prior distribution. More detailed architecture and loss functions can be found in the Method section. Once MpraVAE is trained and model converges, multiple samples can be drawn from *q*(*Z*|*X*,*Y*) and after passing through decoder *θ*, the generated *X̂* can be used for the data augmentation purpose.

**Figure 1.**
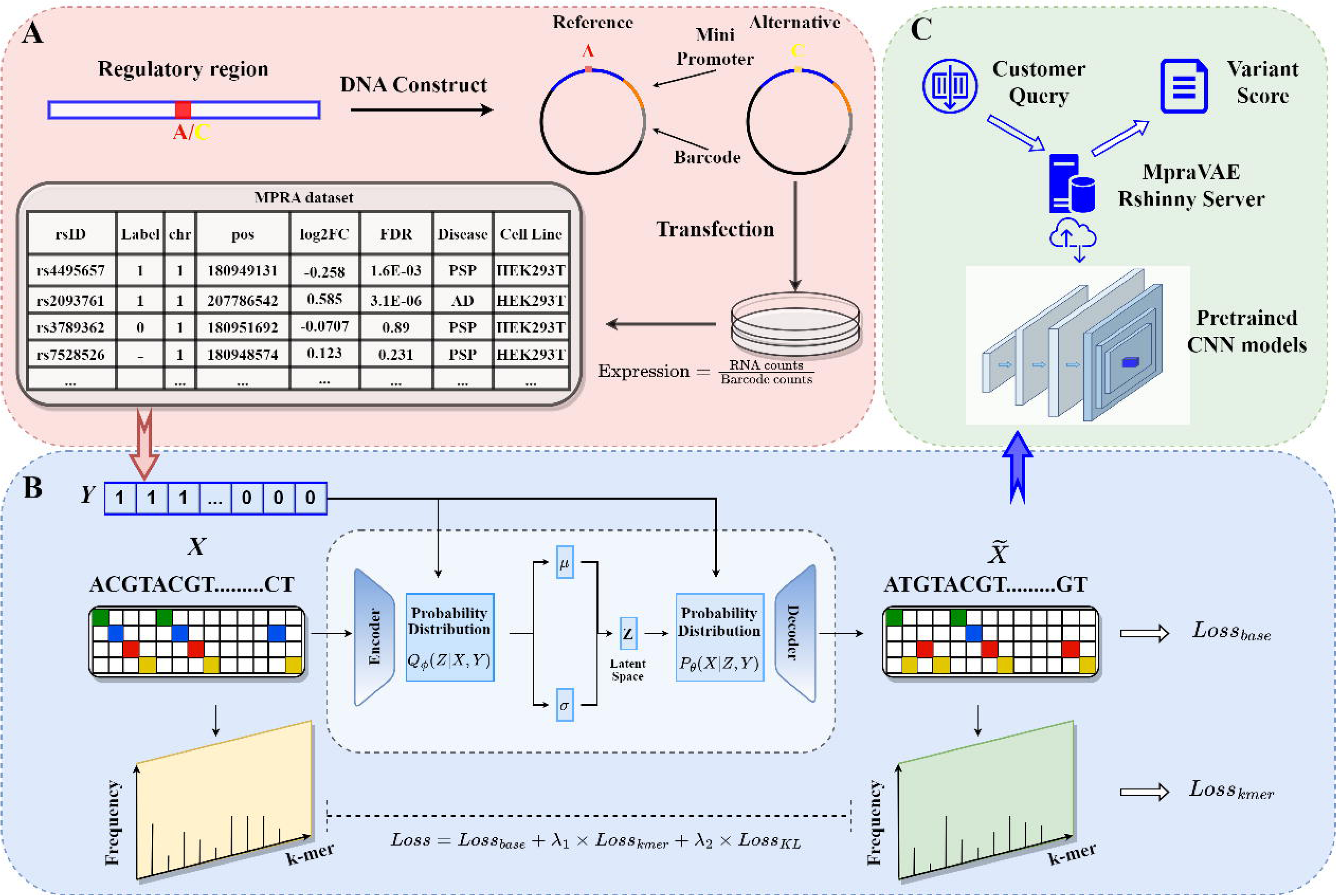
Overview of MpraVAE. **A.** MPRA experiment and data preparation. The labelled variants are derived from MPRA experiments, which focus on evaluating the allelic regulatory effects of variants. Specifically, the MPRA experiment select regulatory sequences that harbor a short polymorphism (SNV or indel), a minimal promoter and a unique DNA barcode that can be transcribed to build the synthetic DNA construct. Two DNA constructs are built for reference and alternative alleles of the variant respectively. These constructs are transfected into cultured cells, and RNA and DNA sequencing are performed to measure the transcription level as the ratio of RNA counts to barcode counts for each allele. To identify variants of significant allelic regulatory effects, transcriptional activity is compared between the two alleles, resulting in a fold change and pvalue. Be leveraging both fold change and pvalue, a binary label can be defined to indicate a positive or negative regulatory variant. The binary label serves as the binary outcome for training data augmentation methods and the CNN classifier. In addition, the surrounding regulatory sequence of the variant, which may regulate the transcription of the barcode sequence, is one-hot encoded as the model input. **B.** Model architecture of MpraVAE. MpraVAE takes the 1000bp one-hot encoding sequences and the binary labels from **A**. as the training set. MpraVAE employs a conditional Variational Autoencoder (CVAE) architecture to generate synthetic sequences that resemble real flanking DNA sequences from positive and negative regulatory to variants, respectively. The encoder learns the conditional latent distribution *Z*, which is a normal distribution, given the sequence *X* and binary label *Y*. The decoder takes both *Z* and *Y* to generate one-hot encoding sequence *X̂*, which can be used for the data augmentation purpose. To learn the network parameters, MpraVAE minimizes the total loss *L_MpraVAE_*, which is defined as the weighted sum of three loss: *L_base_* measuring the difference between real and synthetic sequences on the single base level, *L_kmer_* measuring the difference between real and synthetic sequences on the k-mer level, and the KL-divergence loss *L_KL_* ensuring the learnt latent distribution is close to the prior distribution. Both *L_base_* and *L_kmer_* constitutes the reconstruction loss. **C.** Overview of the MpraVAE web server. The pretrained augmented CNN models, which are trained by both synthetic sequences generated by MpraVAE and real sequences, provide prediction via an RShiny web server. This web server facilitates the usage of MpraVAE by allowing users to input query variants and receive variant scores indicating the probability of being regulatory.

### MpraVAE improves the prediction for regulatory variants in GWAS risk loci compared to existing data augmentation methods

To compare the performance of MpraVAE and existing data augmentation methods for predicting MPRA regulatory variants in GWAS risk loci, we benchmark MpraVAE against the baseline method denoted as “Base” and several simple data augmentation methods denoted as “Base+Rev”, “Base+Crop”, “Base+Rev+Crop”, as adopted in a recent study for improving the prediction for transcription factor binding [31]. Specifically, samples in “Base” include observed flanking sequences of labelled variants identified from MPRA experiments. “Base+Rev” includes samples in “Base” and their reverse complements. For “Base+Crop”, each sequence in “Base” is extended upstream 500bp and downstream 500bp alongside the reference genome, creating two extra sequences by taking the first 1000bp and last 1000bp of the extended sequence. “Base+Rev+Crop” applies this cropping to sequences in “Base” and their reverse complements. These techniques are essentially simple image-based augmentation methods. In addition, we include three variations of semi-supervised methods denoted as “MpraSemi”, “MpraSemi-v1” and “MpraSemi-v2”, which differ in the source of the initial labelled data used for training the CNN classifier and the source of pseudo-labelled data added back. We also compare MpraVAE to MpraVAE-noKmer to evaluate the impact of k-mer loss on the model performance. We evaluate all methods on three MPRA datasets: “MPRA Dementia”, “MPRA autoimmune”, “MPRA melanoma”, which profile variants in strong LD in multiple GWAS loci associated with diseases such as Alzheimer’s disease (AD) and Progressive Suparanuclear Palsy (PSP) autoimmune disease and melanoma. MpraVAE is used to generate synthetic data six times the sample size of “Base”, matching the augmented sample size of "Base+Rev+Crop". For each MPRA dataset, we randomly sample 20% variants as the independent testing set and the remaining 80% as the training set within which 25% variants are randomly sampled as the validation set. To evaluate the impact of training sample size on data augmentation, we construct the augmented training sets using 20%, 50% and 100% of the training set at the baseline. Only scenarios with more than 100 positive variants are evaluated and demonstrated. To ensure a fair comparison, the CNN classifier architecture is kept consistent across all data augmentation methods, while the hyperparameter tuning is performed case by case. Each experiment is repeated 50 times to reduce the bias from random sampling. We plot AUCs and AUPRCs for all methods across different levels of the training set (20%, 50% and 100%). In addition, we use two-sided paired Wilcoxon rank-sum test to evaluate the difference in prediction performance among the methods.

As each study may consist of multiple diseases or cell lines, we evaluate train and evaluate each model stratified by disease or by cell line respectively. For the four diseases under evaluation, the number of positive labelled variants is moderate, ranging from 201 for AD, 410 for PSP, 195 for autoimmune diseases, to 243 for melanoma. When the data is stratified by disease with the 100% training set, we find that MpraVAE consistently outperforms other methods by achieving the largest median of AUCs (mAUC) across four diseases, while the baseline method and other data augmentation methods show significant variation in prediction performance (**Figure 2A**). Specifically, for predicting MPRA regulatory variants associated with AD using labelled data from the “MPRA dementia” study, MpraVAE leads with a mAUC of 0.653, followed by MpraVAE-noKmer and Base+Rev with mAUCs of 0.634 and 0.608 respectively. The improvement in AUC is significant when comparing AUCs from 50 experiments between two methods (pvalue=0.006 for MpraVAE vs. MpraVAE-noKmer; 1.026x10^-5^ for MpraVAE vs. Base+Rev). Similarly, MpraVAE exhibits the largest mAUC of 0.619 for PSP, another dementia disease in the same study. MpraSemi and MpraVAE-noKmer over MpraVAE-noKmer is also significant with a pvalue of 6.935x10^-4^ . In the “MPRA closely follow with mAUCs of 0.617 and 0.608 respectively. The improvement of MpraVAE autoimmune” study, MpraVAE achieves the highest mAUC of 0.732, followed by Base+Crop with 0.710, MpraSemi-v1 with 0.704 and MpraVAE-noKmer with 0.689. Pairwise comparisons between MpraVAE and the three followers show significant improvement (2.577x10^-3^ for MpraVAE vs. Base+Crop; 5.593x10^-3^ for MpraVAE vs. MpraSemi-v1; 2.115x10^-S^ for MpraVAE vs. MpraVAE-noKmer). In the “MPRA melanoma” study, MpraVAE ranks highest with a mAUC of 0.663, followed by MpraVAE-noKmer, MpraSemi-v2 and Base+Rev with mAUCs of 0.657, 0.626 and 0.595, respectively. The pvalues again indicate significant MpraVAE-noKmer; 3.942x10^-6^ for MpraVAE vs. MpraSemi-v2; 2.545x10^-8^ for improvement of MpraVAE compared to its close followers (pvalue=0.034 for MpraVAE vs. MpraVAE vs. Base+Rev). Notably, the baseline model “Base” performs the worst for three out of the four diseases, in contrast to MpraVAE-noKmer, which performs the second best. Meanwhile, three variations of semi-supervised methods and three simple data augmentation approaches demonstrate a large variation in model performance. The superiority of both MpraVAE and MpraVAE-noKmer compared to other approaches demonstrate the effectiveness of using deep learning-based data augmentation approach, specifically, conditional variational autoencoder. In addition, including the loss of k-mer further improves the MpraVAE’s performance, demonstrating the importance to preserve the more spatial correlation within sequences during the data generation. The sample size for Base, unlabeled, and augmented sets using three variations of semi-supervised model to predict four sets of disease-specific regulatory variants are shown in **Supplementary Table S2**.

**Figure 2.**
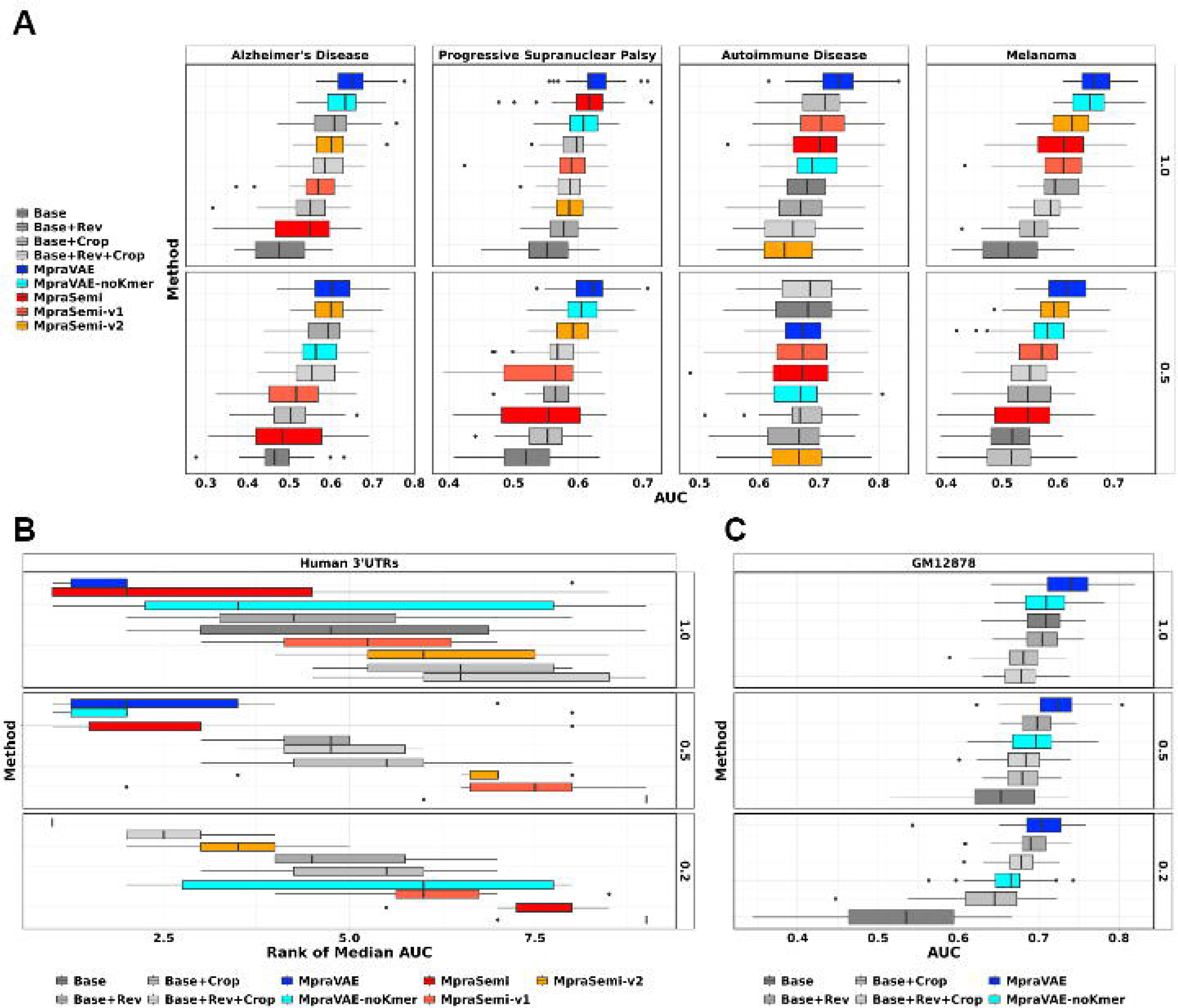
Benchmarking MpraVAE with baseline method, other data augmentation methods and semi-supervised methods in predicting MPRA regulatory variants across five MPRA studies in terms of AUC. **A**. CNN classifiers, which are trained with baseline data and augmented data from eight data augmentation methods on different proportions of training set (50%, 100%), are compared in terms of AUC to predict MPRA labelled variants in GWAS risk loci across four diseases. **B.** CNN classifiers, which are trained with baseline data and augmented data from eight data augmentation methods on different proportions of training set (20%, 50%, 100%) are compared in terms of AUC rank per cell type to predict MPRA variants in human 3’UTRs across six cell types. **C.** CNN classifiers, which are trained with baseline data and augmented data from eight data augmentation methods on different proportions of training set (20%, 50%, 100%) are compared in terms of AUC to predict MPRA validated variants in eQTL loci. For the baseline method, all samples are partitioned into training, validation, and testing sets at a ratio of 60%:20%:20%. The 20% testing set will be used for both baseline method and all data augmentation methods. Data augmentation method is applied to the remaining 80% data to generate synthetic data. For data augmented approaches, the union of real data and synthetic data are split into the training and validation sets at a ratio of 75%:25%. To reduce the bias from random partition, we repeat the random partitions 50 times and report the distribution of AUCs.

We further downsample the training set to 50% to evaluate the impact of training sample size on the model performance of data augmentation approaches. As expected, the overall performance of all methods decrease compared to 100% training set, indicating the training sample size impacts on the performance of data augmentation methods. The richness of training samples contributes to the robustness of both baseline model and the augmented model for the unseen prediction. Despite the decreasing of the baseline samples, MpraVAE achieves the best overall performance except for autoimmune disease, where all methods demonstrate competitive performance. Taken together, these observations demonstrate that MpraVAE can improve the prediction of disease-specific regulatory variants compared to both baseline model and other data augmentation methods when the training size is reasonably large. Moreover, similar trends are observed when the prediction performance is measured by AUPRC (**Figure S1**).

### MpraVAE improves the prediction for regulatory variants in functional genomic regions

Besides being an invaluable tool to evaluate the regulatory potential of variants in strong LD within GWAS loci associated with phenotypes such as diseases or traits, MPRA can also been employed to localize individual causal variants influencing molecular phenotypic traits (e.g., eQTL) and to target specific functional genomic regions (e.g., 3’UTR) due to their unique functional roles. For example, 3′UTRs contain a particularly important class of noncoding variants that play crucial roles in post-transcriptional and translational processes. Accordingly, we apply MpraVAE and other data augmentation approaches to two MPRA studies: “MPRA 3’UTR” and “MPRA GM12878”. “MPRA 3’UTR” extensively evaluates variants in 3′ untranslated regions (3′UTRs) in six human cell lines. “MPRA GM12878” evaluates variants associated with 3,642 eQTLs in lymphoblastoid cell lines to identify variants that directly modulate gene expression. The training, validation testing sets are constructed in the same way as before, with each experiment repeated 50 times. The training set is downsized to different proportions (100%, 50% and 20%) to evaluate its impact on model performance. As each variant in the “MPRA 3’UTR” study is evaluated in six human cell lines, we train each model in a cell line-specific manner and obtain the AUC/AURPC for each method for each cell line. For each cell line, we rank the AUC/ AUPRC for all compared methods and report the distribution of AUC/AUPRC ranks and median of AUC/AUPRC ranks (mAUCrank/mAUPRCrank).

When using the full training set for data augmentation, MpraVAE leads the prediction performance by achieving the highest rank with a mAUCrank of 2, on par with MpraSemi. It is followed by MpraVAE-noKmer with a mAUCrank of 3.5 and Base+Rev with 4.25 (**Figure 2B**). However, MpraVAE exhibits a much lower variability of ranks across cell lines (7.067) than both MpraSemi (9.175) and MpraVAE-noKmer (12.267). Using AURPC as the metric, MpraVAE demonstrates both a higher mAUPRCrank of 1 and lower variability of 0.700 than MpraSemi (mAUPRCrank=3; variability=2.800) and MpraVAE-noKmer (mAUPRCrank=2; variability=13.067). Specifically, MpraVAE ranks 1st in 2 cell lines (HMEC, HEPG2) and 2nd in 3 cell lines (HEK293FT, K562, SKNSH) measured by AUC and 1st in 4 cell lines (HEK293FT, HEPG2, K562, SKNSH) and 2nd in 1 cell line (GM12878) measured by AURPC.

At the 50% training size, MpraVAE and MpraVAE-noKmer and MpraSemi show comparable performance with mAUCranks of 2, 2 and 3 respectively. Nevertheless, MpraVAE achieves the highest mAUPRCrank of 1, with variability 1.467, compared to MpraVAE-noKmer and MpraSemi, both with a mAUPRCrank of 2.5 (variability=10.667 and variability=1.467, respectively). Notably, the baseline model performs the worst, with a mAUCrank of 9 and mAUPRCrank of 9, indicating the importance of data augmentation in improving the prediction performance, especially when the baseline sample size is small. Similar trends are observed with 20% training set, where MpraVAE remains the top-performing method. Some differences are also observed. MpraVAE performs the best across all six cell lines, while the baseline model performs the worst in terms of both mAUCrank and mAUPRCrank. In addition, the performance of both MpraVAE-noKmer and MpraSemi is significantly deteriorated. Overall, MpraVAE demonstrates a clear advantage over the baseline method and existing data augmentation methods in predicting regulatory variants in 3’UTR.

As unlabeled data is missing in the processed “MPRA GM12878” study, we exclude semi-supervised methods from the comparison. MpraVAE outperforms other methods by MpraVAE-noKmer (AUC=0.710, pvalue= 3.113x10^-4^) and baseline (AUC=0.709, achieving highest mAUC of 0.740 with the 100% training set, significantly higher than pvalue=8.361x10^-S^) (**Figure 2C**). When the training set is reduced to 50% and further to 20%, MpraVAE still leads all methods by obtaining the highest mAUC of 0.723 and 0.705 respectively. In both scenarios, MpraVAE significantly outperforms the secondary-ranked Base+Rev, which has a mAUC of 0.698 (pvalue=4.105x10^-4)^ and 0.690 (pvalue=9.675x 10^-3)^. Moreover, the baseline model performs worst with the 50% and 20% training sets, further demonstrating the advantage of augmenting the training set through data augmentation approaches. In addition, the performance of MpraVAE-noKmer is significantly deteriorated as the training sample size decreases. All methods exhibit a similar trend in performance when AUPRC is used as the metric (**Figure S1B,C**).

### MpraVAE outperforms existing variant scoring methods

By demonstrating MpraVAE as the top-performing data augmentation method, we further compare MpraVAE-augmented CNN classifier to a list of existing variant scoring approaches, which encompass a wide range of supervised and unsupervised methods, leveraging their pre-computed functional scores, on 3 MPRA datasets targeting GWAS loci and 2 MPRA datasets targeting functional genomic regions. The same data split strategy is adopted for creating training and testing sets, and 50 repeated experiments are performed to avoid sampling bias. Performance is measured by both AUC and AUPRC using the full training set, followed by a two-sided paired Wilcoxon rank-sum test to assess AUC/AUPRC differences for pairwise method comparison.

We find that MpraVAE consistently outperforms other methods in terms of AUC across four diseases **(Figure 3A)**. Additionally, other approaches show a large variability of performance. Specifically, for AD, MpraVAE leads with a mAUC of 0.653, followed by FIRE with 0.575, and the improvement is significant with a pvalue of 1.542x10^-8^. For PSP, another dementia disease, MpraVAE achieves the highest mAUC of 0.619, which significantly (pvalue=9.930x10^-10^). Interestingly, Orion performs the worst for both dementia diseases, outperforms the second-ranked FATHMM-MKL that obtains a mAUC of 0.536 with mAUCs of 0.487 and 0.486 respectively, which is close to a random guess. For autoimmune diseases, MpraVAE achieves the highest mAUC of 0.732, significantly higher than the next method, FitCons2, with a mAUC of 0.642 (pvalue= 9.622x10^-S^). Similarly, for melanoma, FitCons2, with a mAUC of 0.562. The pvalue of this pairwise comparison is 1.055x10^-9^, MpraVAE demonstrates the highest mAUC of 0.663, surpassing the subsequent method, indicating significant superiority of MpraVAE over FitCons2. Interestingly, we find that FIRE, which performs best for AD, performs worst for Melanoma. FATHMM-MKL, the top-ranked existing method for PSP, ranks close to bottom for both AD and Melanoma. In addition, Orion consistently has poor performance across all four diseases. Despite the variable performance of other methods, MpraVAE consistently ranks at the top, demonstrating its robust performance. The same evaluation is performed on “MPRA 3’UTR” and “MPRA GM12878”. For MPRA 3’UTR, the ranks of AUC for each method across all six cell types are reported. MpraVAE achieves a mAUCrank of 1 and mAUPRCrank of 1, which significantly outperforms the succeeding method, FATHMM.XF, with a mAUCrank of 2.5 and mAUPRCrank of 2.0 (**Figure 3B, S2B)**. For MPRA GM12878, MpraVAE leads the prediction performance with a mAUC of 0.740 and mAUPRC of 0.411, followed by DVAR with a mAUC of 0.662 and FunSeq2 with a mAUPRC of 0.386 (**Figure 3C, S2C)**.

**Figure 3.**
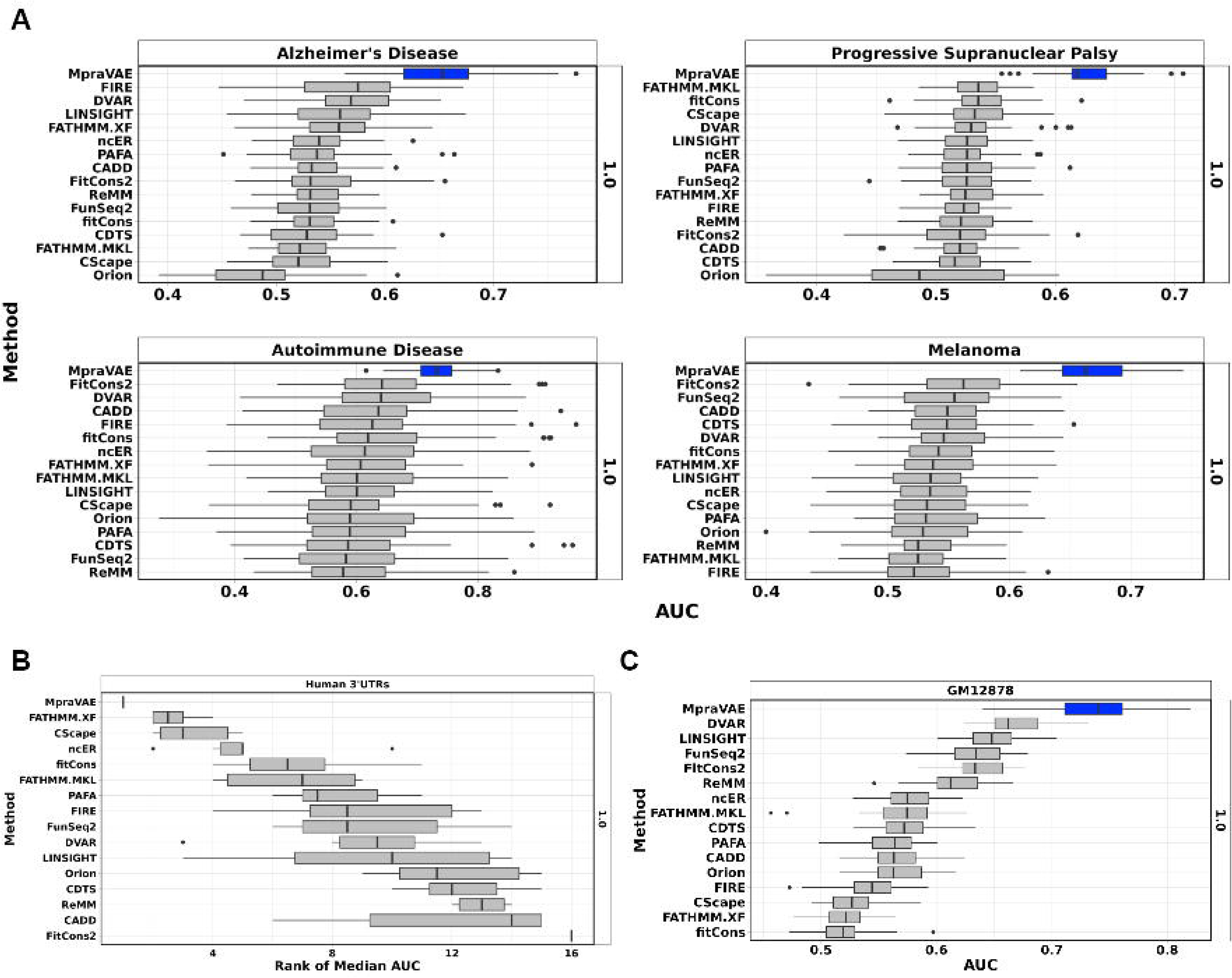
Benchmarking MpraVAE-augmented classifier and 15 variant scoring methods in predicting MPRA regulatory variants across five MPRA studies in terms of AUC. 20% of both positive and negative variants are randomly sampled to form an independent testing set. Data augmentation method is applied to the remaining 80% data to generate synthetic data. For data augmented approaches, the union of real data and synthetic data are split into the training and validation sets with a ratio of 75%:25%. To reduce the bias from random partition, we repeat the random partitions 50 times and report the distribution of AUCs.

### Genome-wide characterization of MPRA regulatory activities associated with autoimmune diseases

Using the labelled variants from MPRA studies, MpraVAE-augmented classifier demonstrates superior performance compared to existing methods in aforementioned sessions. A nature extension of the application of the pretrained MpraVAE-augmented classifier, is to perform a genome-wide screen to obtain MPRA regulatory activities and investigate their global patterns. Without loss of generality, we adopt the MpraVAE-augmented classifier, which is trained by the labelled variants from the “MPRA autoimmune” study, to obtain a genome-wide functional score for all SNVs on hg19 from 1000 Genomes provided by CADD [32]. To differentiate high regulatory variants from the background ones, we define positive variants as those ranking in the top 95th percentile of the functional score, while background variants encompass the remaining variants.

As MPRA experiments aim to assay the regulatory effects of individual variants on gene expression, we are interested in assessing the distribution of the positive and background variants in the cis-regulatory elements (CREs), which are known for regulating the transcription of genes. Particularly, enhancers, a crucial type of CRE, control gene expression program, which is mediated by chromatin looping that brings enhancers close to promoters. In addition, enhancers also exhibit tissue- and cell type-specific activities, so the evaluation will be performed in a tissue- and cell type-specific way [33]. Accordingly, we will evaluate the enrichment pattern of MPRA variants in cell type-specific enhancers. Specifically, we collect cell type-specific human enhancers from EnhancerAtlas 2.0 and narrow down to nine cell types related to the immune system [34]. The enrichment is calculated as the proportion of variants overlapped with cell type-specific enhancers among all variants for the positive set and background set respectively. Consequently, we observe the enrichment of positive set is consistently higher than that of the background set across the nine cell types, indicating MPRA variants are more likely to play a functional role by locating within enhancers **(Figure 4A**).

**Figure 4.**
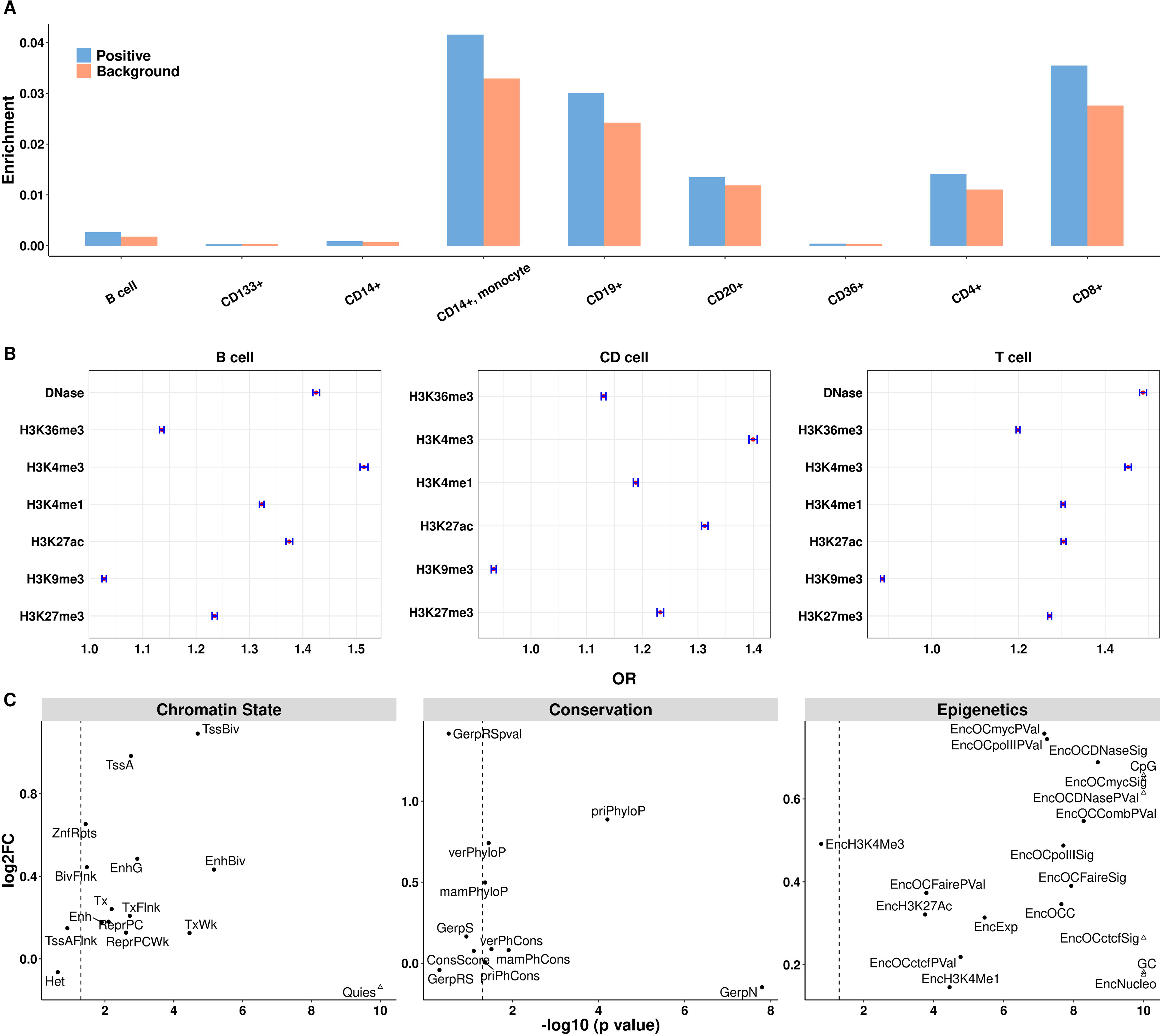
Genome-wide characterization of MPRA regulatory variants associated with autoimmune diseases. **A.** The prevalence of positive and background variants defined by MpraVAE in immune cell type-specific enhancer regions from EnhancerAtlas 2.0. The enrichment of positive and background variants defined by MpraVAE in open chromatin region and histone modification site across B cell, CD cell and T cell, which are measured by odds ratio (red) and its 95% confidence interval (blue). **C.** Comparing MpraVAE positive and background variants across three categories of functional annotations including chromatin state, conservation score and epigenetics. log2FC indicates the log2 of the ratio of mean annotation value between positive and background variants. -log10(p-value) represents -log10 of pvalue from testing the difference of mean annotation value between positive and background variants. Dashed line represents -log10 (0.05). A blank triangle represents a pvalue less than 10^−10^.

Next, we investigate the enrichment of MPRA variants in genome-wide histone modification as their activities are closely associated with gene expression. We collect ChIP-seq peaks from B cell, T cell and CD cell, which are three immune-related cell types, across multiple histone marks and chromatin accessibility from Roadmap Epigenomics Project. We then calculate the odds ratio (OR) as the ratio between number of positive variants in peaks/number of positive variants not in peaks and number of background variants in peaks/number of background variants not in peaks. We also report the 95% confidence interval of each OR. Consequently, we find that positive variants are overall more enriched in active histone marks and open chromatin than background variants (**Figure 4A,B**). The highest enrichment has been observed for open chromatin and H3K4me3, a chromatin mark associated with promoter activation. Specifically, positive variants are enriched in open chromatin regions, measured by DNase-seq, of both B cell and T cell with ORs of 1.425 (1.419,1.431) and 1.487 (1.479,1.495) respectively. They are also enriched in H3K4me3 of all three cell types with ORs of 1.515, 1.400 and 1.453. A similar trend of enrichment can be observed for chromatin marks associated with enhancer activation such as H3K4me1 (1.323, 1.188, 1.303), H3K27ac (1.375, 1.312, 1.304) and chromatin marks associated with gene expression activation such as H3K36me3 (1.135, 1.130, 1.199) across three cell types. All the above observations indicate that the predicted regulatory variants are more likely located in open chromatin regions, enhancer and promoters to have a functional impact. Interestingly, two repressive histone marks show opposite enrichment patterns. Positive variants are enriched in H3K27me3, while no enrichment is observed for H3K9me3. One possible explanation is that H3K27me3-rich regions can be used to identify silencers that can regulate gene expression via proximity or looping and the regulatory variants may exert their function via silencers [35]. Overall, though the results are expected and not supervising, the findings underscore our model’s efficacy in identifying regulatory variants enriched in CREs in a cell type-specific way.

Moreover, we evaluate the distribution difference of three categories of CADD functional annotations between the positive and background variants, which include chromatin state, conservation score and epigenetics. To facilitate the analysis, we randomly select 2,000 positive variants and 40,000 background variants, ensuring proportional representation for both variant sets in each chromosome, and we calculate the log2 fold change between the two sets for each category of functional annotation and report the pvalue using the Wilcoxon sum-rank test. For chromatin state, we find that most chromatin states are more enriched in the positive set than the background set with fold change larger than 1 and pvalue less than 0.05. Specifically, four (log2FC=0.983, pvalue=1.750x10^-3^), an active transcription start site-proximal promoter state chromatin states demonstrate a large effect size and significant difference. These include TssA associated with expressed genes; enhancer state EnhG (log2FC=0.485, pvalue1.146x10^-3^); TSSBiv (log2FC=1.091, pvalue=2.017x10^-S^) and EnhBiv (log2FC=0.433, pvalue=6.800x 10^-6^), which are two inactive states consist of bivalent regulatory states of promoter and enhancer respectively. Nevertheless, most conservation scores are less pronounced contrast between positive and background sets, with most of the pvalues around 0.05. priPhyloP (Primate size (log2FC=0.887, pvalue=6.244x10^-S^), while GerpN (Neutral evolution score defined by Phylo Pscore) is the only conservation score showing both significant difference and large effect 0.148, pvalue=1.600x10^-8^). Notably, most of functional scores in the epigenetics show much GERP++) only shows statistically significant difference but the effect size is small (log2FC=-higher enrichment in the positive set. Examples include epigenetic annotations related to proteins that transcribes gene such as EncOCmycSig, EncOCmycPVal (Peak signal and pvalue for Myc evidence of open chromatin) and EncOCpolIISig, EncOCpolIIPVal (Peak signal and pvalue for Pol II evidence of open chromatin). Other epigenetic annotations include open chromatin regions profiled by DNase-seq and FAIRE-seq such as EncOCDNaseSig, EncOCDNasePVal (Peak signal and pvalue for DNase evidence of open chromatin) and EncOCFaireSig, EncOCFairePVal (Peak signal and pvalue for Faire evidence of open chromatin), as well as combined evidence of CREs such as EncOCCombPVal (combined pvalue of Faire, DNase, PolII, CTCF, Myc evidence for open chromatin). In addition, epigenetic annotations related to active histone mark, DNA methylation and gene expression, which include EncH3K27Ac (Maximum ENCODE H3K27 acetylation level), CpG (Percent CpG in a window of +/- 75bp), EncExp (Maximum ENCODE expression value), also show significant enrichment in the positive set. These observations align with the expectation that predicted MPRA regulatory variants are intricately linked to CREs and other key genomic elements that regulate the gene expression.

### Linking MpreVAE predicted regulatory variants associated with autoimmune diseases to target genes by integrating multi-omics data

Leveraging cell type-specific epigenomic data and PCHi-C (Promoter Capture Hi-C) from immune-related cell types, we can link the predicted regulatory variants to target genes. Identifying these target genes may reveal how these variants play a functional role in disease, potentially by affecting the expression of the target genes. Moreover, the SNP-to-gene link can help validate that the predicted variants are causal, especially if the targeted genes have known biological function related to the disease or trait. Specifically, we collect DNase-seq tracks in CD3, CD4, CD8, Naïve CD4 and Th1 cell types from ENCODE and PCHi-C data from the same cell types [36]. For example, we spotlight one variant rs2815403 (chr10:103893336), which exhibits a high MpraVAE score of 0.683 and located in the intergenic region of PPRC1 gene (chr10:103892787-103910090) (**Figure 5A**). In addition, rs2815403 resides in an open chromatin region consistent across all five immune cell types, indicated by DNase signals. rs2815403 is further linked by a PCHi-C chromatin loop to another open chromatin region consistent across all five immune cell types in the promoter region of NFKB2, a gene that is part of the NFkB complex, which functions as a central activator of genes involved in inflammation and immune response by regulating cytokine and chemokine expression [37, 38]. Another variant rs11719339 (chr3:27410561) achieves a remarkably high MpraVAE score of 0.993 and is also found in an open chromatin region consistent across all five immune cell types, as indicated by DNase signals, within the NEK10 (chr3:27152394-27410912). Linked by a chromatin loop inferred from PCHi-C, rs11719339 interacts with the promoter region of STAT2, a member of the STAT family that plays an essential role in immune responses to extracellular and intracellular stimuli [39]. The above observation confirms that DNase I hypersensitive sites (DHSs) are a hallmark of chromatin regions containing CREs such as promoters. In addition, disease-associated regulatory variants are enriched in close proximity to DHSs, which may disrupt transcription factor binding sites and contribute to disease by altering the expression of putative target genes identified through chromatin loop via the SNP-to-gene link [40]. Moreover, as both targeted genes are immune-related, the SNP-to-gene link further validates the two highly scored variants are causal. In short, the above examples demonstrate the effectiveness of the deep learning model enhanced by MpraVAE in identifying regulatory variants and enable the discovery of their regulated disease genes by integrating with multi-omics data such as HiC and ChIP-seq.

**Figure 5.**
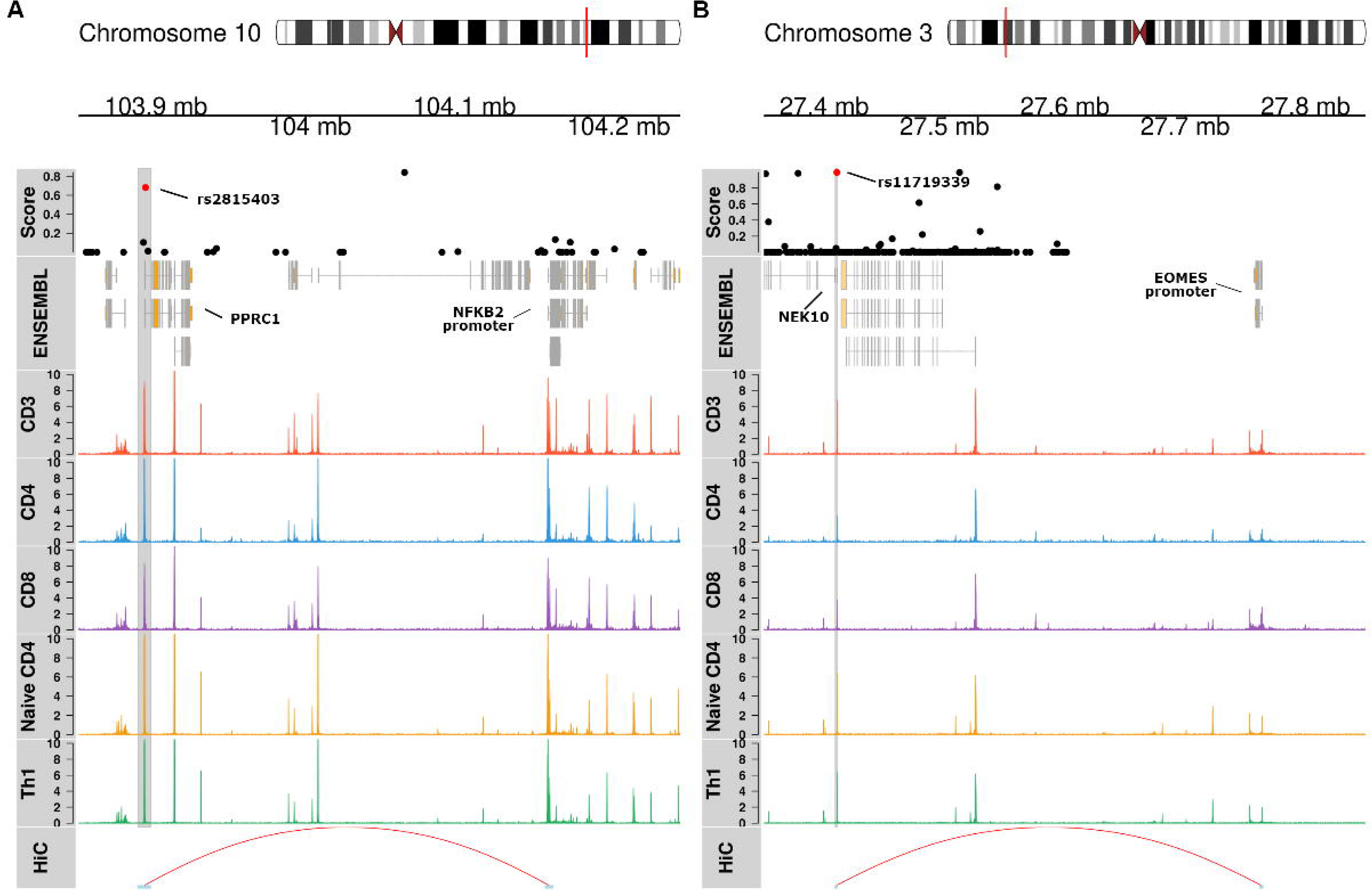
Linking MpreVAE predicted regulatory variants associated with autoimmune diseases to target genes by integrating multi-omics data. The score track displays the predicted regulatory scores for variants in linkage disequilibrium (LD) (R>0.2) with the targeted variant (red). The ENSEMBL track presents gene coordinates from Ensembl GRCh37. The following epigenetic tracks demonstrate the peaks representing open chromatin regions derived from DNase-seq in immune cell types. The final “HiC” track shows a chromatin loop derived from promoter-capture HiC (PCHi-C) data, which is present across all immune cell types. The chromatin loop anchors on the region containing the variant of interest and another region of gene promoter. Two novel variants and targeted immune-related genes are identified. **A.** rs2815403 and NFKB2 **B.** rs11719339 and EOMES.

## Discussion

In this work, we present MpraVAE, a deep generative model, designed to augment the sample size of labelled variants generated from MPRA experiment for improving the prediction performance for regulatory variants. Instead of generating the labelled variants directly, MpraVAE leverages the one-hot encoding flanking DNA sequence surrounding the labelled variants and augment the sequence representation. As the MPRA variants are usually labelled as positive and negative according to the strength of allelic regulatory effect, we develop MpraVAE in the framework of conditional variational autoencoder and thus generating the positive and negative labelled variants respectively. One significant contribution of MpraVAE is the construction of the reconstruction loss at both the base and k-mer levels to improve the structure similarity between synthetic and real sequences. By benchmarking MpraVAE on five MPRA datasets, which target variants in GWAS loci, eQTL loci and 3’UTR, we demonstrate that MpraVAE outperforms simple data augmentation approaches and semi-supervised learning methods that leverage the unlabeled variants. Notably, MpraVAE performs better than its closest counterpart, which excludes k-mer loss in the reconstruction loss. Additionally, MpraVAE-augmented classifier surpasses a wide range of supervised and unsupervised variant scoring methods.

We further extend the application of MpraVAE by performing a genome-wide screen to obtain MPRA regulatory activities and investigate their global patterns by using autoimmune diseases as one example. We find that positive variants are enriched in enhancers, open chromatin regions, active histone marks associated with promoter, enhancer and gene body in immune-related cell types. They are also more frequently present in chromatin states associated with promoter, enhancer and binding sites of cMyC and Pol II that regulate gene expression. These observations demonstrate that MPRA regulatory variants are more likely to reside in the active cis-regulatory elements to exert their impact on the gene expression. By leveraging cell type-specific epigenomic data and PCHi-C from immune-related cell types, we link the predicted MPRA regulatory variants to target genes by chromatin loops. Through two examples, we successfully link highly scored MRPA variants to their targeted genes, which are found to play a crucial role in the immune response. As both targeted genes are immune-related, the SNP-to-gene link further validates the two highly scored variants are causal. In addition, both variants and promoter of targeted genes are enriched of DNase signals. These variants are enriched in close proximity to DHSs, which may disrupt transcription factor binding sites and contribute to disease by altering the expression of target genes identified through chromatin loop via the SNP-to-gene link. Lastly, we develop a RShinny web server, hosting the pretrained MpraVAE models, the facilitates the usage of MpraVAE by allowing the users to upload variants and obtain the prediction score.

Several possible extensions of MpraVAE can be made to further improve the model performance. Currently, we only employ the DNA flanking sequence in the neighborhoods of labelled MPRA variants as the feature for data augmentation. In addition to DNA sequence, functional annotations surrounding the DNA flanking sequence can be also integrated in the data augmentation process by developing a multi-modal conditional variational autoencoder framework. Specifically, multi-track of functional annotations of tissue/cell type-specific omics data, such as histone modification, transcription factors, can be stacked in multi-dimension matrix, with entry values obtained by calculating signals on the binned DNA sequence. The feasibility is supported by the distribution difference of functional annotations, such as enhancer, histone marks and chromatin states, between positive and background variants in the studies of the autoimmune diseases. Moreover, we focus on performing data augmentation on MPRA labelled variants. With the decreasing of sequencing cost, the application of MpraVAE can be readily adapted to labelled variants generated from other experimental approaches such as CRISPR. A comprehensive evaluation of the feature and label extension can be pursued in the future work.

## Methods and Materials

### MPRA studies

We collect the summary data from five Massively Parallel Reporter Assay (MPRA) studies targeting GWAS loci, eQTL loci and 3’UTR. These studies specifically screen noncoding variants in tight LD within GWAS and eQTL loci as well as in key genomic regions such as 3’UTR to identify the causal variants. For each study, variants showing allelic regulatory effects with a False Discovery Rate (FDR) less than 0.1 and log2 fold change greater than 0.1 are deemed as positive regulatory variants. Conversely, variants with FDR larger than 0.8 are labeled as “negative variants”. The remaining variants are categorized as “unlabeled”. To ensure stable training and testing, we only consider the experiments that have more than 100 positive variants. The five MPRA studies are as follows: (i) “MPRA Dementia”: This study focus on Alzheimer’s disease (AD) and Progressive supranuclear palsy (PSP), which are two dementia diseases.

MPRA experiment is performed to screen noncoding SNVs in 25 significant GWAS loci associated with AD and nine loci associated with PSP in the HEK293T cell line [41]. The dataset includes 201 positive, 655 negative and 882 unlabeled variants for AD, and 410 positive, 1352 negative and 1840 unlabeled variants for PSP. (ii) “MPRA autoimmune”: This study screens variants in tight LD (r2>0.8) with 578 GWAS index variants representing 531 distinct GWAS loci associated with five Autoimmune diseases: Type 1 diabetes (T1D), Ulcerative colitis (UC), Crohn’s disease (CD), Rheumatoid arthritis (RA), Multiple sclerosis (MS) in Jurkat T cells [42]. The dataset consists of 195 positive, 15,960 negative and 2,150 unlabeled variants. To address the substantial imbalance between numbers of positive and negative variants, 1950 negative variants are sampled to maintain the 10:1 ratio of negative to positive variants. (iii) “MPRA melanoma”: This study functionally screens risk variants in tight LD (r2>0.8) from 54 loci associated with melanoma [43]. (iv) “MPRA 3’UTR”: This study performs a genome-wide functional screen of 3′UTR variants associated with human disease and evolution. The 3′UTR variants are selected from those in LD from the NHGRI-EBI GWAS catalog, overlapping regions identified as being under positive selection in humans and rare 3’UTR variants in genes with outlier expression signatures across tissues in GTEx. The 3′UTR variants are profiled in six cell lines (HMEC, HEK293FT, HEPG2, K562, GM12878, SKNSH) [44]. Consequently, the positive set ranges 405 to 1188 variants, negative set ranges 4207 to 8361 variants and unlabeled set ranges 3311 to 6703 variants across six cell lines. (v) “MPRA GM12878”: This study screens variants associated with eQTLs in lymphoblastoid cell lines. The labelled variants are selected by He et al. with 693 positive controls and 2772 negative controls [45].

### Model architecture for MpraVAE

To perform data augmentation on labelled variants, MpraVAE leverages the variant representation in the form of one-hot encoding flanking sequence surrounding labelled variants, essentially a 2D image to facilitate the usage of the deep generative model. Specifically, each sequence has the variant as mid-point extending 500 bp upstream and downstream, as the model input. The sequence is further one-hot encoded as A: [1,0,0,0], C: [0,1,0,0], G: [0,0,1,0] and T: [0,0,0,1]. MpraVAE employs a Conditional Variational Autoencoder (CVAE) architecture to generate synthetic sequences that resemble flanking DNA sequences surrounding labelled MPRA variants [46, 47]. MpraVAE consists of a conditional encoder *φ* and a conditional decoder *θ*. The encoder *φ* takes both the one-hot encoding sequence *X*, representing the variant, and the label for the variant *Y* (1 for positive and 0 for negative) as input to learn the conditional latent distribution *q*(*Z*|*X*,*Y*) of the latent variable *Z* given the sequence *X* and label *Y*. *q*(*Z*|*X*,*Y*) is essentially a normal distribution *N*(*μ*(*X*,*Y*),*σ*^2^(X,*Y*)) with parameters are estimated from both *X* and *Y*. The decoder takes both *Z* and *Y* to generate the reconstructed one-hot encoding sequence denoted as *X̂*. Specifically, the encoder *φ* takes one-hot encoding sequences and the binary class label *Y* of MPRA variant (1 or 0) as the input, which directly connects to a convolutional layer. Each convolutional layer is followed by a one-dimensional batch normalization layer and activated by the Rectified Linear Unit (ReLU). This block of convolutional and batch normalization layers is repeated twice. The output of the last normalization layer is connected to two fully connected layers to obtain the parameters *μ* and *σ*^2^.

The decoder *θ* takes latent variable *Z*, drawn from *N*(*μ*(*X*,*Y*),*σ*^2^(*X*,*Y*)), and the class label as the input. This is followed by a fully connected layer and a batch normalization layer, which is activated by ReLU. The output is reshaped into a 3D tensor and passed through a transposed convolutional layer, followed by another normalization layer and ReLU activation. The output is then connected to a second transposed convolutional layer, activated by the softmax function, to generate the probability of one-hot encoding sequences.

To learn the network parameters, MpraVAE minimizes the total loss *L_mpraVAE_*, which is defined as the weighted sum of three loss,

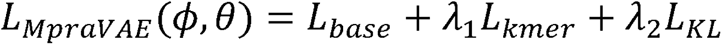

Specifically, *L_base_* measures the difference of between real and synthetic sequences on the single base level, which is formularized as,

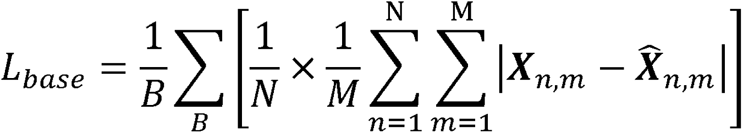

where *B* is the batch size, *N* the flanking sequence length, *M* is the number of bases, ***X*** and ***X̂*** represents one real sequence and one synthetic sequence respectively. *L_kmer_* measures the difference between real and synthetic sequences on the k-mer level, which is formularized as,

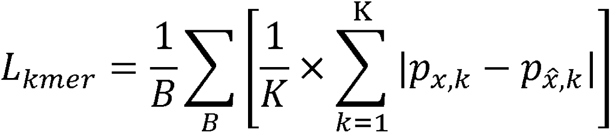

where k-mer is defined a substring of length *k* within the flanking sequence, *P_x,k_* and *P_x,k_* represent the proportion of k-mer frequency in ***X*** and ***X*** respectively. *K* is the number of k-mer combinations. Without loss of generality, we set *k* as 3 since it will produce a non-sparse representation of k-mer proportion and *K* is therefore 64 for all 3mer combinations. The introduction of *L_kmer_* enhances the measurement of similarity between real and synthetic sequences, recognizing that nucleotides are spatially correlated to form important genomic element such as motifs. The reconstruction loss comprises both *L_base_* and *L_kmer_* to reflect the similarity between synthetic and real sequences on different levels of resolution. The third loss *L_kL_* is the KL-divergence loss *L_KL_* (q(*Z*|*X*,*Y*)||*P*(*Z*)), which will ensure the learnt latent distribution is close to the prior distribution. In addition, weight parameters *λ*_1_ and *λ*_2_ are applied to *L_kmer_* and *L_KL_* to adjust the magnitude of each individual loss, ensuring each loss impacts the model training process. Once MpraVAE is trained and model converges, multiple samples can be drawn from q(*Z*|*X*,*Y*) and after passing through decoder *θ*, the generated *X̂* can be used for the data augmentation purpose.

### Convolutional neural network as the classifier

Given either the original or augmented dataset, a classifier is needed to predict a given variant is regulatory. Considering the nature of the variant representation, which is one-hot encoding sequence, and inspired by the previous works, we adopt the convolutional neural network (CNN) as the classifier [48–51]. Particularly, The CNN takes one-hot encoding sequences as the input, which directly connects to a convolutional layer. Each convolutional layer is activated by ReLU and followed by a max pooling layer and a dropout layer. The block of convolutional-max pooling-dropout layer is repeated three times. The output of last dropout layer is flattened and connected to three dense layers. Each dense layer is activated by ReLU and followed by a dropout layer, enabling hidden neurons in each dense layer randomly dropped out to avoid overfitting. The output of last dropout layer is connected to the output layer, which is activated by the softmax function to predict the probability of a given variant being regulatory.

### Data augmentation methods

To evaluate the effectiveness of MpraVAE, we benchmark it against a series of data augmentation methods. The first set of three approaches are inspired by a study that augments DNA sequences to improve the DNA-protein binding prediction [52]. Specifically, the original DNA sequences surrounding MPRA labelled variants are deemed as the baseline named “Base”. Accordingly, a CNN classifier trained using samples from “Base” is considered as the baseline method. “Base+Rev” includes all sequences in “Base” and their reverse complements, resulting in a sample size twice that of “Base”. For “Base+Crop”, Each sequence in “Base” is further extended upstream 500bp and downstream 500bp alongside the reference genome. Two extra sequences are created by taking the first 1000bp and last 1000bp of the extended sequence.

Therefore, “Base+Crop” has a triple sample size that of “Base”. “Base+Rev+Crop” applies this cropping to sequences in “Base” and their reverse complements, resulting in a sample size six times that of “Base”. Additionally, to assess the effect of considering *L_kmer_* in the reconstruction loss, we include "MpraVAE-noKmer", which removes *L_kmer_* from the total loss during the model training. By default, both MpraVAE and MpraVAE-noKmer are trained using sequences in “Base” and will generate synthetic data that is six times the size of “Base”. For all data augmented approaches, the labels of augmented sequences are treated the same as their corresponding original sequences.

### Semi-supervised learning

Semi-supervised learning is an approach that improves the prediction performance by leveraging both labeled and unlabeled data, in contrast to traditional supervised learning techniques that rely solely on labeled data. Here, we introduced a semi-supervised learning algorithm named "MpraSemi" to augment the training set by leveraging both labelled and unlabeled MPRA variants [53]. Unlike data augmentation methods, which generate synthetic training samples using only labelled MPRA variants, MpraSemi boosts the training set by incorporating additional samples from unlabeled MPRA variants. The working flow of MpraSemi is illustrated in **Figure S3**. All MPRA variants are classified into three categories “positive”, “negative” and “unlabeled” based on the log2FC and FDR cutoff. For each iteration of the algorithm, only positive and negative labeled variants serve as the input for the CNN classifier, which predicts the probabilities of being positive for both labeled and unlabeled data, and the predicted probability distribution are denoted as *P̂_Labeled_* and *P̂_unlabeled_* respectively. We then obtain the 10th and 90th percentiles from, *P̂_Labeled_* denoted as *P̂*_10_ and *̂P*_90_. Unlabeled data with predicted probabilities lower than *P̂*_10_ are labelled as “pseudo negative”, while those higher than *P̂*_90_ are labelled as “pseudo positive”. The remaining samples are retained as "unlabeled". These pseudo-labeled samples are then integrated into the labeled data for subsequent iterations. This iterative process continues until either no new pseudo samples are identified or a predefined threshold for the iteration loops are reached. Moreover, we introduce three variations of MpraSemi based on the source of the initial labelled data used for training the CNN classifier and the source of pseudo-labelled data added back, denoted as “MpraSemi”, “MpraSemi-v1” and “MpraSemi-v2” respectively. MpraSemi adopts samples from "Base" as the initial input for both training CNN classifier and generating pseudo-labeled samples. MpraSemi-v1 uses samples from "Base+Rev+Crop" as the initial labelled data for training the CNN classifier but only utilizes samples from "Base" for generating pseudo-labeled samples. MpraSemi-v2 extends the usage of samples from "Base+Rev+Crop" for both the initial training CNN classifier and generating pseudo-labelled data. The variations of MpraSemi help evaluate the combined effect of the semi-supervised learning and naïve data augmentation approach in predicting regulatory variants.

### Model training, validation and testing

All deep learning models are implemented using PyTorch on an NVIDIA A100 GPU system [54]. Utilizing mini-batch gradient descent and the Adam optimizer, the CNN network is optimized for binary outcome using cross-entropy loss [55]. The default learning rate is set to 10^-3^. To improve the efficiency of learning process, warm-up steps and a learning rate decay strategy are optionally incorporated. Each model undergoes training for a maximum of 200 epochs, with early stopping implemented if the model performance stagnated over 10 consecutive epochs. Considering the inherent heterogeneity across the MPRA datasets, the hyperparameters of the CNN models are fine-tuned independently for each disease/cell type-specific model. The encoder-decoder architecture of MpraVAE is trained in a similar way.

For the baseline method, all samples from “Base” are partitioned into training, validation, and testing sets at a ratio of 60%:20%:20%. The 20% testing set will be used for both baseline method and all data augmentation methods. Data augmentation method is applied to the remaining 80% data to generate synthetic data. For data augmented approaches, the union of observed data and synthetic data are split into the training and validation sets with a ratio of 75%:25%. The prediction performance of baseline and data augmentation-driven CNN classifier is measured by the Area Under the Receiver Operating Characteristic Curve (AUC) and the Area Under the Precision-Recall Curve (AUPRC). To reduce the bias from random partition, we repeat the random partition 50 times and compare the AUC or AUPRC using paired Wilcox sum-rank test. To perform a sensitivity analysis to evaluate the impact of baseline sample size on the prediction performance, we down-sample the original labelled variants to 20% and 50% fractions, while ensuring the positive set contains at least 100 variants. We then repeat the same experiments.

### Variant scoring methods

We adopt a wide collection of both supervised and unsupervised methods for variant scoring as a comparison for MpraVAE. For unsupervised methods, we include fitCons and FitCons2, which are evolutionary model to assess the fitness consequences of point mutations [9, 10]; FunSeq2, a weighted scoring method that integrates conservation score, transcription factor binding events, enhancer-gene linkages etc. [8]; Orion, an approach that detects regions of non-coding genome that are depleted of variation [56]; CDTS, a method that uses the absolute difference of the observed variation from the expected variation [57]; and DVAR, an approach that leverages biochemical and evolutionary evidence to assess functional impact of noncoding variants [11].

Regulatory variants in Human Gene Mutation Database (HGMD), pathogenetic variants in ClinVar, variants in NHGRI GWAS Catalog (https://www.ebi.ac.uk/gwas/) and somatic mutations in COSMIC have been widely used as labelled variants for training various supervised methods [12–14]. For example, both FATHMM-XF and FATHMM-MKL uses HGMD [15, 16]; PAFA adopts ClinVar and GWAS SNPs [58]; CScape utilizes COSMIC [18]; and ncER [17] employs both ClinVar and HGMD. Other methods such as CADD [59] adopts high-frequency human-derived alleles; ReMM uses curated regulatory variants involved in Mendelian disease [60]; LINSIGHT utilizes SNPs in 54 unrelated individuals from Complete Genomics [61]. These methods also vary in terms of used prediction models: FATHMM-XF, FATHMM-MKL, CADD adopt SVM; ncER, FIRE and ReMM use tree-based machine learning methods such as XGBoost and random forest, while LINSIGHT and PAFA utilize Generalized Linear Model.

## Code and Data availability

MpraVAE is available at github with the link (https://github.com/yi-xiaa/MpraVAE). The R Shinny web server of MpraVAE is available at https://mpravae.rc.ufl.edu/

## Acknowledgements

This work was supported by National Institute of General Medical Sciences of the National Institutes of Health under Award Number R35GM142701 to LC.

## Author contributions

L.C. conceived and designed the study. WJ.J. and Y.X. and SR.T. performed the experiments. WJ.J., Y.X., L.C. wrote the manuscript with input from all the other authors. All authors have read and approved the final manuscript.

## Ethics declarations

The authors declare no competing interests.

## Supplementary information

**Table S1. Labelled, unlabeled and increased sample size by semi-supervised model when stratified by disease for the five MPRA studies**

**Table S2. Labelled, unlabeled and increased sample size by semi-supervised model when stratified by cell line for the five MPRA studies**

**Figure S1. Benchmarking MpraVAE with baseline method, other data augmentation methods and semi-supervised methods in predicting MPRA regulatory variants across five MPRA studies in terms of AURPC. A**. CNN classifiers, which are trained with baseline data and augmented data from eight data augmentation methods on different proportions of training set (50%, 100%), are compared in terms of AURPC to predict MPRA labelled variants in GWAS risk loci across four diseases. **B.** CNN classifiers, which are trained with baseline data and augmented data from eight data augmentation methods on different proportions of training set (20%, 50%, 100%) are compared in terms of AURPC rank per cell type to predict MPRA variants in human 3’UTRs across six cell types. **C.** CNN classifiers, which are trained with baseline data and augmented data from eight data augmentation methods on different proportions of training set (20%, 50%, 100%) are compared in terms of AURPC to predict MPRA validated variants in eQTL loci. For the baseline method, all samples are partitioned into training, validation, and testing sets at a ratio of 60%:20%:20%. The 20% testing set will be used for both baseline method and all data augmentation methods. Data augmentation method is applied to the remaining 80% data to generate synthetic data. For data augmented approaches, the union of real data and synthetic data are split into the training and validation sets at a ratio of 75%:25%. To reduce the bias from random partition, we repeat the random partitions 50 times and report the distribution of AURPCs.

**Figure S2. Benchmarking MpraVAE-augmented classifier and 15 variant scoring methods in predicting MPRA regulatory variants across five MPRA studies in terms of AUPRC.** 20% of both positive and negative variants are randomly sampled to form an independent testing set. Data augmentation method is applied to the remaining 80% data to generate synthetic data. For data augmented approaches, the union of real data and synthetic data are split into the training and validation sets with a ratio of 75%:25%. To reduce the bias from random partition, we repeat the random partitions 50 times and report the distribution of AUPRCs.

**Figure S3. Overview of MpraSemi.** All MPRA variants are classified into three categories “positive”, “negative” and “unlabeled”. For each iteration of the algorithm, only positive and negative labeled variants serve as the input for training a CNN classifier. The trained CNN classifier predicts the probabilities of being positive for both labeled and unlabeled data. for labeled data (*p̂*_10_) are labelled as “pseudo negative”, while those higher than 90th percentile Unlabeled data with predicted probabilities lower than 10th percentile of predicted probabilities of predicted probabilities for labeled data (*p̂*_90_) are labelled as “pseudo positive”. The remaining samples are retained as "unlabeled". Subsequently, these pseudo-labeled samples are integrated into the labeled data for subsequent iterations. This process is repeated until no more pseudo-labels can be assigned or a preset number of iterations is completed.

