## Supplementary materials for "In silico generation and augmentation of regulatory variants from massively parallel reporter assay using conditional variational autoencoder"

**Supplementary tables**

**Table S1. Labelled, unlabeled and increased sample size by semi-supervised model when stratified by disease for the four MPRA studies**

| **Disease** | **Labeled Sample Size** | | **Unlabeled Sample Size** | **Increased Sample Size (MpraSemi/MpraSemi_v1)** | **Increased Sample Size (MpraSemi_v2)** |
| --- | --- | --- | --- | --- | --- |
|  | **Pos** | **Neg** |  |  |  |
| **Functional regulatory variants implicate distinct transcriptional networks in dementia** | | | | | |
| AD | 201 | 655 | 882 | 530 | 332 |
| PSP | 410 | 1352 | 1840 | 818 | 604 |
| **Prioritization of autoimmune disease-associated genetic variants that perturb regulatory element activity in T cells** | | | | | |
| Autoimmune Disease | 195 | 1950 | 16156 | 4217 | 4965 |
| **Massively parallel reporter assays and variant scoring identified functional variants and target genes for melanoma loci and highlighted cell-type specificity** | | | | | |
| Melanoma | 243 | 1109 | 2628 | 954 | 1332 |
| **A semi-supervised approach for predicting cell-type specific functional consequences of non-coding variation using MPRAs** | | | | | |
| GM12878 | 693 | 2772 | N/A | N/A | N/A |

**Table S2. Labelled, unlabeled and increased sample size by semi-supervised model when stratified by cell line for the five MPRA studies**

| **Cell line** | **Labeled Sample Size** | | **Unlabeled Sample Size** | **Increased Sample Size (MpraSemi/MpraSemi_v1)** | **Increased Sample Size (MpraSemi_v2)** |
| --- | --- | --- | --- | --- | --- |
|  | Pos | Neg |  |  |  |
| **Functional regulatory variants implicate distinct transcriptional networks in dementia** | | | | | |
| HEK293T | 611 | 2007 | 2722 | 987 | 882 |
| **Genome-wide functional screen of 3′UTR variants uncovers causal variants for human disease and evolution** | | | | | |
| GM12878 | 548 | 6932 | 4618 | 1373 | 1333 |
| HEK293FT | 572 | 8112 | 3414 | 1092 | 821 |
| HEPG2 | 426 | 8361 | 3311 | 1158 | 806 |
| HMEC | 1188 | 4207 | 6703 | 1513 | 2761 |
| K562 | 410 | 8297 | 3391 | 1109 | 1018 |
| SKNSH | 405 | 7764 | 3929 | 1351 | 1012 |
| **Prioritization of autoimmune disease-associated genetic variants that perturb regulatory element activity in T cells** | | | | | |
| Jurkat | 195 | 1950 | 16156 | 4217 | 4965 |
| **Massively parallel reporter assays and variant scoring identified functional variants and target genes for melanoma loci and highlighted cell-type specificity** | | | | | |
| Malignant Melanoma | 176 | 668 | 1146 | 564 | 422 |
| Normal Melanocyte | 67 | 441 | 1482 | 606 | 714 |
| **A semi-supervised approach for predicting cell-type specific functional consequences of non-coding variation using MPRAs** | | | | | |
| GM12878 | 693 | 2772 | N/A | N/A | N/A |

**Figure S1.** **Benchmarking MpraVAE with baseline method, other data augmentation methods and semi-supervised methods in predicting MPRA regulatory variants across five MPRA studies in terms of AURPC.**

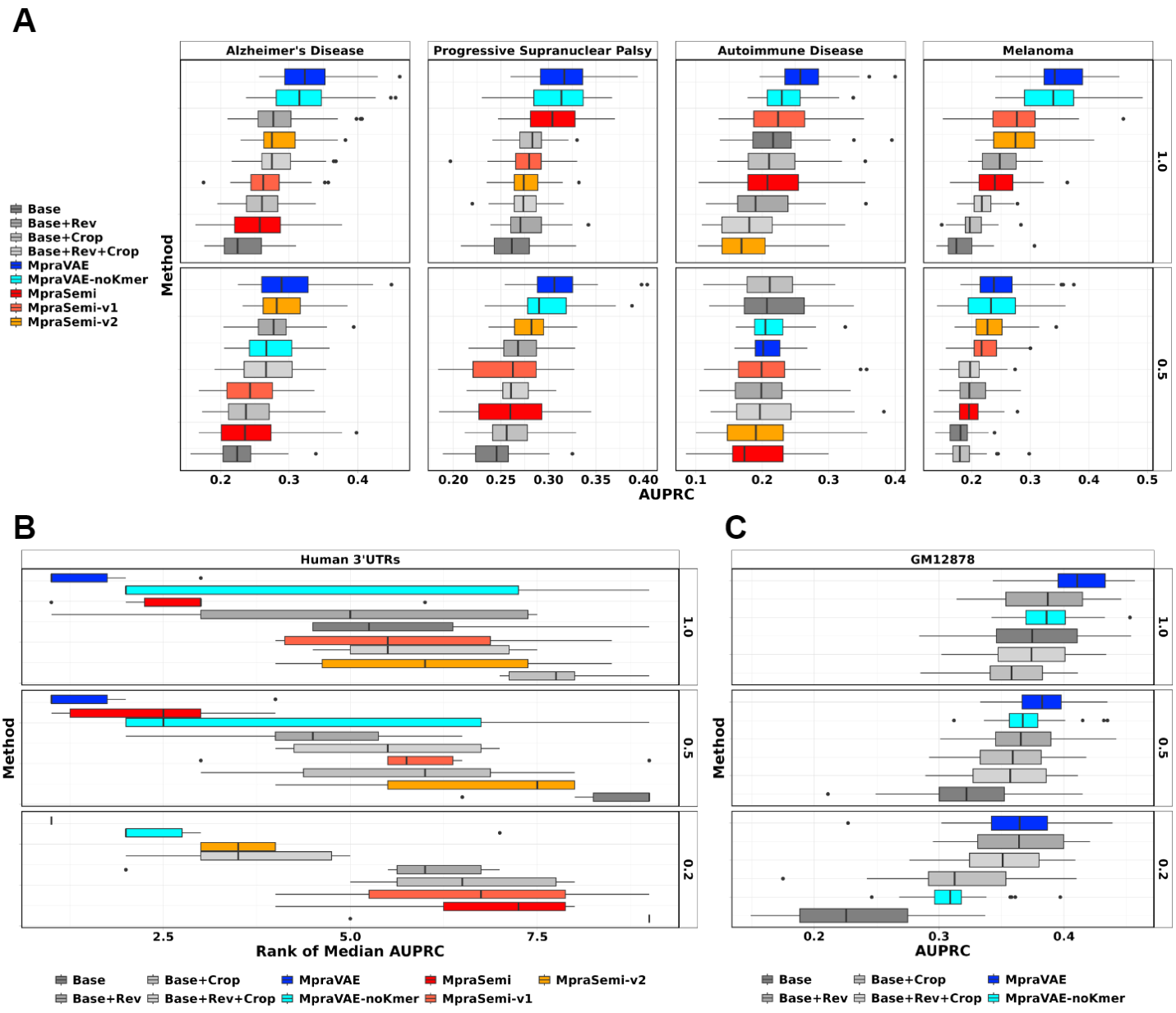

**Figure S2.** **Benchmarking MpraVAE-augmented classifier and 15 variant scoring methods in predicting MPRA regulatory variants across five MPRA studies in terms of AUPRC.**

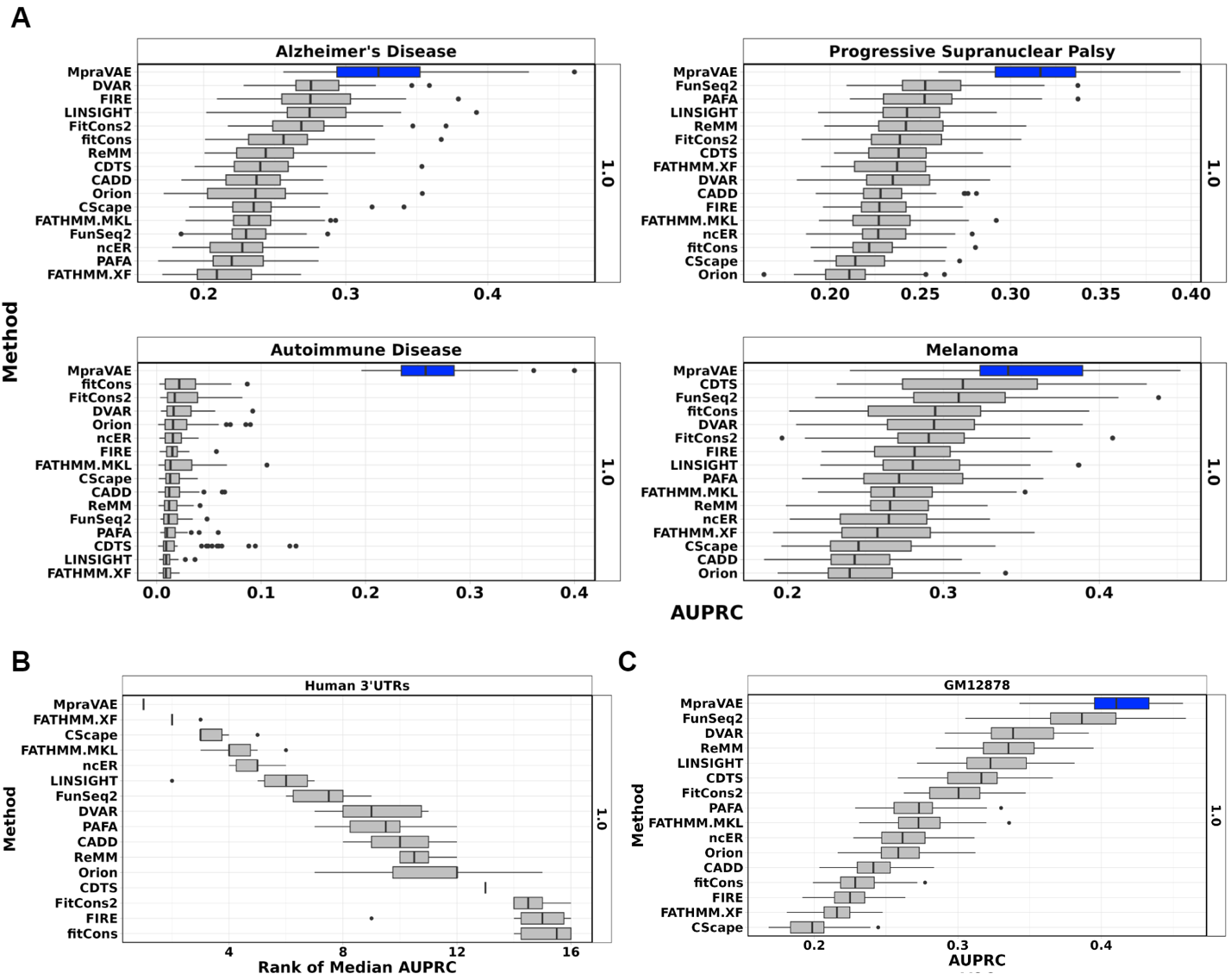

**Figure S3.** **Overview of MpraSemi.**
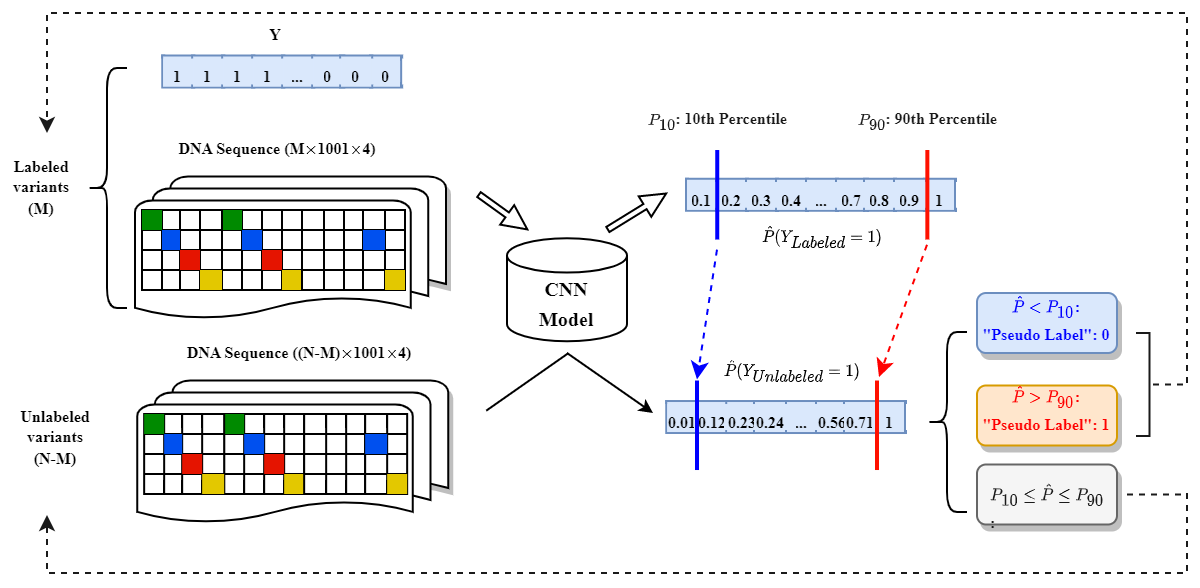
